# Comprehensive online two-dimensional nanoLCxCZE-MS for deep top-down proteomics

**DOI:** 10.64898/2026.05.14.725123

**Authors:** Tobias Waldmann, Philipp T. Kaulich, Andreas Tholey, Christian Neusüß

**Affiliations:** Faculty of Chemistry, Aalen University, 73430 Aalen, Germany; Systematic Proteome Research & Bioanalytics, Institute for Experimental Medicine, Christian-Albrechts-Universität zu Kiel, 24105 Kiel, Germany

**Author notes:** Correspondence: Prof. Dr. Christian Neusüß, Beethovenstr. 1, 73430 Aalen, Germany.

## Abstract

Understanding proteoforms, i.e., the various molecular forms in which proteins can exist, is important for deciphering biological processes and diseases. While capillary zone electrophoresis (CZE) proved advantageous for proteoform separation, limited sample loading capabilities restrict its application. Here, we present a novel comprehensive two-dimensional nanoLCxCZE-MS platform for deep top-down proteomics (TDP). The 2D platform is highly automated, enabling robust performance and the possibility to perform proteoform quantitation as demonstrated by isobaric labeling experiments. The high orthogonality of reversed-phase LC and CZE leads to a peak capacity of 2200, leading to an increase in the number of identified proteoforms in a human Caucasian colon adenocarcinoma cell lysate sample by a factor of 3 compared to nanoLC-MS. Furthermore, CZE mobilities enable the attribution of many more proteoforms to a certain proteoform family on the MS1-level. Overall, the flexible platform enables highly efficient separation of intact proteoforms combined with sensitive MS-based TDP workflows, both for untargeted and targeted analysis of complex biological samples.

**Graphical Abstract:** We report a robust and automated comprehensive nanoLCxCZE-MS platform for top–down proteomics. In addition to large volume sample injection and separation by hydrophobicity in the nanoLC, the orthogonal separation by CZE in the second dimension leads to a strong increase in peak capacity and, thus, in the number of identified proteoforms. CZE mobilities also enable the attribution of many more proteoforms to a proteoform family on the MS1-level.

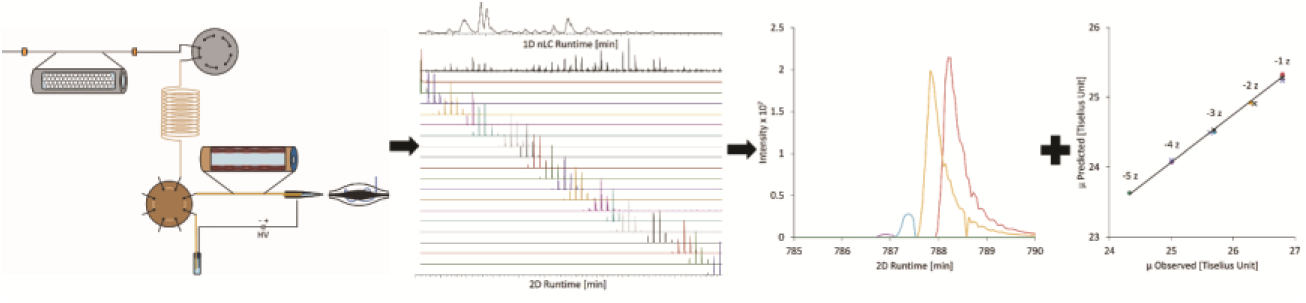

## Introduction

The lower-than-expected number of genes identified by the human genome project led to the conclusion that biological complexity is rather on the protein level than on the genomic level ^[1]^. Therefore, analysis of proteins is crucial to understanding biological systems. Complexity on the protein level not only arises from genetic variation or alternative splicing but also from posttranslational modification (PTM) of proteins. All those processes result in a plethora of distinct molecules, called proteoforms, that modulate a wide variety of biological processes ^[1,2]^. A group of related proteoforms forms a proteoform family ^[3]^.

Proteoform-level analysis by top-down proteomics (TDP) has great potential for identifying key diagnostic markers as well as therapeutic targets. Identification and quantification of proteins at the proteoform level are thus essential for researching human disease phenotypes ^[4]^. However, intact proteoform analysis is challenging, e.g., due to front-end separation of intact proteins. While reversed-phase liquid chromatography (RPLC) is the predominant method, capillary zone electrophoresis (CZE) emerges as a promising alternative ^[5,6]^. CZE-MS can be especially useful to separate intact proteoforms within a proteoform family varying in their electrophoretic mobility (charge-to-size ratio), e.g., due to phosphorylation or acetylation ^[7,8]^. Nevertheless, one-dimensional separation techniques are often not sufficient to adequately separate intact proteoforms in complex samples ^[5,9,10]^. To increase proteome coverage, multidimensional separation techniques are typically used ^[5,11]^. Gel-eluted liquid fraction entrapment electrophoresis (GELFrEE), as well as the more recently introduced PEPPI (passively eluting protein from polyacrylamide gel as intact species), are popular pre-fractionation techniques ^[5,12,13]^. Size-based pre-fractionation techniques, such as size exclusion chromatography, provide a detergent-free alternative ^[5,10]^. However, offline protein fractionation requires large sample quantities due to sample loss during sample collection and transfer, has low throughput, and is labor-intensive ^[5,10,11]^. Online coupled multidimensional techniques, such as 2D high-pH RPLC/low-pH RPLC, typically offer higher sensitivity, as well as potentially reduced sample degradation due to minimized and automated sample handling between the dimensions ^[14]^. In addition to 2D LC, coupling of RPLC with CZE seems promising, since it allows for large-volume sample injection in the first dimension while separating proteoforms based on an orthogonal separation mechanism in the second dimension (hydrophobicity and electrophoretic mobility, respectively). Early approaches in online LC-CZE-MS coupling date back to the 1990s and involved either loop-based ^[15,16]^ approaches or flow-gating interfaces ^[17,18]^. Although valve-based approaches that physically separate the two dimensions are advantageous, e.g., for capillary coating and flushing ^[19]^, nowadays, a majority of online LC-CZE coupling uses flow gating approaches on microfluidic chips ^[20–23]^. Only a few studies coupled online HPLC and CZE for TDP ^[24]^. Recently, we reported online coupling of reversed-phase nanoLC and CZE-MS for heart-cut and targeted proteoform analysis on the intact protein level using an insulated Polyether ether ketone (PEEK) valve with transfer volumes of 20 nL ^[25,26]^. In an initial attempt, we were able to transfer a certain fraction of the LC stepwise into the CZE ^[26]^.

Nevertheless, for deep top-down proteoform characterization, a holistic approach is desirable, in which all separated proteins and proteoforms from the first-dimensional nanoLC separation are transferred to the second dimension (comprehensive 2D). However, to the best of our knowledge, no comprehensive online nanoLCxCZE-MS approach for TDP has been described so far.

Here, we report an automated, comprehensive nanoLCxCZE-MS platform for intact proteoform characterization and proteoform family assignment by TDP. The setup allows storing the separated proteoforms from the first-dimensional nanoLC separation (several µL), and automatically transferring them to the second dimension for further CZE-MS characterization. This enables us to perform, for the first time, a comprehensive, in-depth, untargeted proteome-wide nanoLCxCZE-MS analysis of complex biological samples, demonstrated here using a human Caucasian colon adenocarcinoma (CaCo-2) cell lysate. Due to the additional selectivity in the second dimension (separation according to ion mobilities), the platform has great potential in proteoform characterization, especially within a proteoform family. The method was evaluated against 1D nanoLC regarding orthogonality, number of proteins/proteoforms identified, and the influence of sample concentration. Additionally, we performed iodoTMT-labeled quantitative analysis and demonstrated that proteoform mobility can serve as a powerful piece of information to validate identified proteoforms or assign proteoforms to proteoform families at the MS1-level.

## Results and Discussion

### Setup of the NanoLCxCZE-MS Platform

Prior to mass spectrometric characterization, proteoform separation is important to reduce co-elution and overlapping signals and obtain high-quality MS and MS/MS information. Therefore, we developed an automated, comprehensive, online coupled two-dimensional nanoLCxCZE-MS platform. First, the proteoforms are separated based on hydrophobicity using reversed-phase nanoLC. Subsequently, the proteoforms are separated in the second dimension based on their electrophoretic mobility in the liquid phase using capillary zone electrophoresis, which is online coupled via nanoESI to top-down mass spectrometry.

For a typical online comprehensive coupling in the liquid phase, two small loops are alternately sampled and injected into a very fast second-dimensional separation capable of analyzing the sample at the same time as the next fraction is collected in the other sample loop ^[27]^. Due to the small injection volumes in CZE, such a setup is not feasible with commercial CE systems, where typical separations are in the mid-minute range. Here, we present a setup capable of comprehensive nanoLCxCZE-MS with time-decoupling of the two separation dimensions by storing the first-dimension effluent in a capillary and subsequently transferring one fraction after another into the CZE. A sketch of the setup and its principal operation is shown in Figure 1.

**Figure 1:**
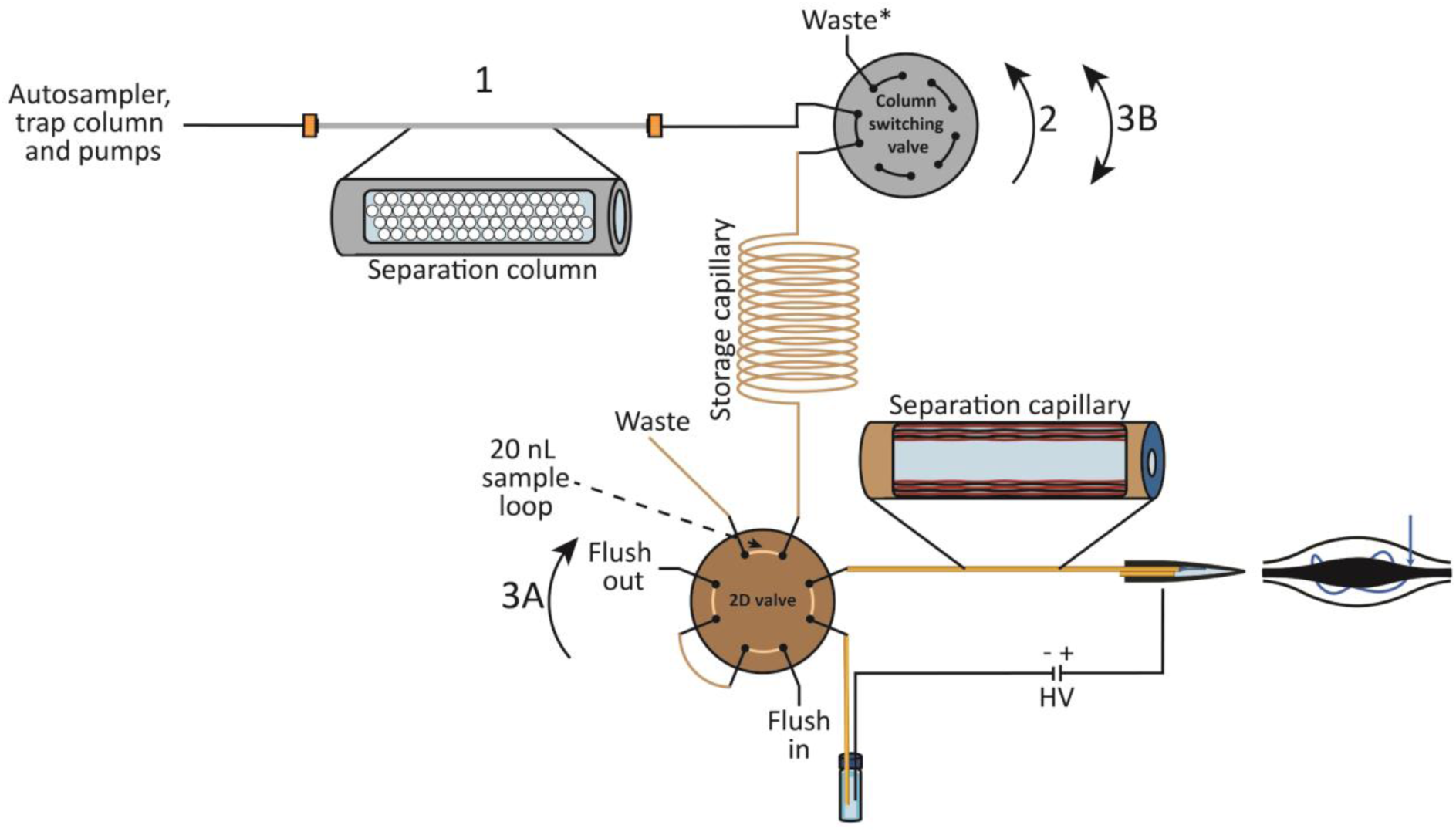
Workflow of the nanoLCxCZE-MS platform. (1) Separation of the sample on the reversed-phase column (nano LC). (2) Decoupling of the storage capillary from the nanoLC-flow as soon as a fraction of interest is inside the storage capillary. (3A) Upon switching the 2D valve, the content of the 2D valve sample loop (20 nL) is separated in the second dimension. (3B) The next 2D valve sample loop is filled with the stored fraction by coupling and decoupling of the storage capillary with the nanoLC flow for a defined period of time (e.g., 0.6 min at a flow rate of 100 nL/min to deliver 60 nL to the storage capillary). Steps 3A and 3B are repeated until the stored fraction is completely transferred to the second dimension. Unused ports and sample loops of the 2D valve are flushed with water. *The waste capillary can also be connected to a nanoLC-MS interface to perform 1D nanoLC-MS measurements.

In our nanoLCxCZE-MS platform, the sample is injected using a “trap-and-elute” approach to enable large volume injection (up to 20 µL) followed by nanoLC separation, where the effluent is guided through the column-switching valve into the storage capillary with a user-defined volume for subsequent analysis by CZE-MS (Figure 1, step 1). When the fraction to be characterized in the second dimension (e.g., the whole protein-containing fraction of the first dimension for a comprehensive approach) is in the storage capillary, the nanoLC column is decoupled using the column-switching valve (Figure 1, step 2). Sample transfer from the storage capillary into the connected 2D valve for injection into the CZE-MS is carried out by switching the column-switching valve in a fixed time interval for a defined time, thus coupling and decoupling the storage capillary with the nanoLC flow. For the here applied nanoLC flow rate of 100 nL/min, switching for 0.2 min transfers 20 nL from the storage capillary through the sample loop of the 2D valve (Figure 1, step 3B). This process is later on referred to as “pulsing” of a “pulsed volume”. This 2D valve is a customized commercial Polyether ether ketone (PEEK) valve with four internal sample loops to transfer 20 nL fractions to the second dimension. This PEEK 2D valve can be used for subsequent online CZE-MS analysis because the loops are compatible with the injection volume of CZE-MS analysis, and the valve material (PEEK) is not conductive and insulates the fluidic path. Thus, a high electrical field can be applied through the valve for the subsequent CZE separation. The procedure of switching the column-switching valve for a defined time (Figure 1, step 3B), followed by switching of the 2D valve (Figure 1, step 3A) and subsequent CZE separation, is repeated until the stored content is completely analyzed by CZE-MS (comprehensive mode). Sample storage, transfer, and subsequent analysis in the second dimension are fully automated and universal, enabling various applications depending on available measurement time and the desired coverage of the first dimension. Thus, the platform can, e.g., be used for targeted applications in a heart-cut or selective comprehensive approach (Figure S 1 A-C), or for untargeted applications in a comprehensive approach (Figure S 1 D). For the latter, a large storage volume is required to store all nanoLC-separated analytes. Here, we use a storage capillary with a capacity of 3.5 µL (180 cm x 50 µm ID), which enables the storage of 35 min of the nanoLC separation.

Furthermore, the highly flexible setup allows 1D nanoLC-MS measurements by keeping the column-switching valve permanently connected to the nanoViper capillary (waste in Figure 1), while connecting the nanoViper capillary to an MS by a nanoESI interface. Also, 1D CZE-MS can be performed using the CE autosampler for sample injection, which we used for QC experiments with model proteins (Myoglobin, Ribonuclease A, and Lysozyme). 1D CZE-MS reference measurements of the CaCo-2 sample were performed without the 2D Valve.

### Comprehensive nanoLCxCZE-MS

Applying the method improvements highlighted in the experimental section of the supporting information, an overall two-dimensional separation of the sample was carried out in 11 h (70 min nanoLC, followed by ∼590 min to sample the stored volume). The resulting separation is shown in Figure 2 A.

**Figure 2:**
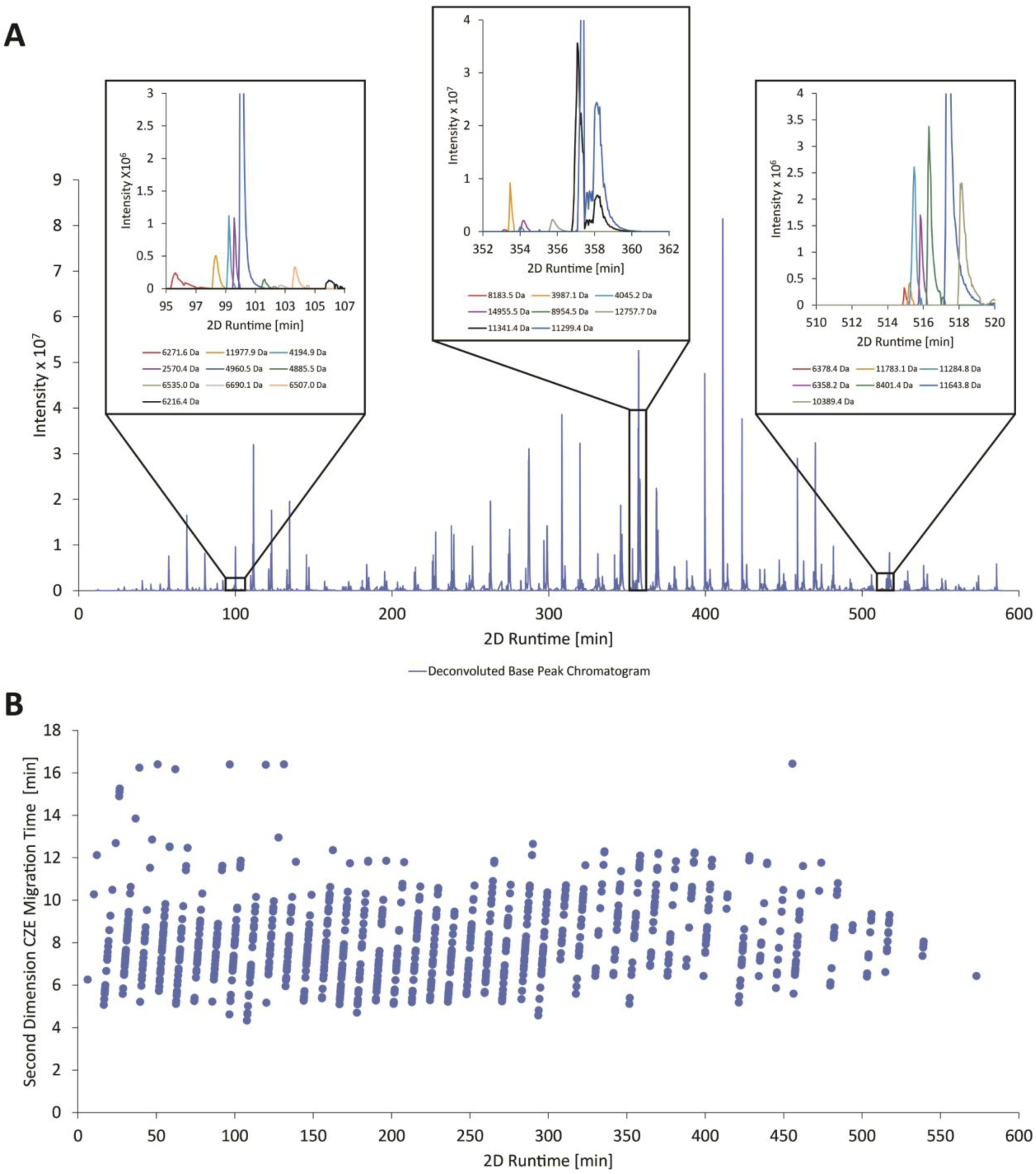
Orthogonality of CZE and nanoLC. A) Base peak electropherogram after deconvolution of a nanoLCxCZE-MS run (blue line) with three zoomed regions randomly selected from the beginning, middle, and end of the 2D measurement. The zoomed regions highlight the high orthogonality of the used nanoLC and CZE dimensions. B) Migration times in the second dimension versus total 2D runtime. Migration times in the second dimension were calculated by linking the reported 2D runtime by ProSightPD (grouped processing of the undiluted sample) to the time the valve was switched for that cut. Next, the two times were subtracted, resulting in the migration time of the proteoform in the second dimension (n=1013). Since the EOF is detected at ∼5 min, Proteoforms with a calculated migration time less than 5 minutes (n=79) were assumed to be proteoforms belonging to the previous cut, except if those proteoforms had a predicted charge of zero (n=7) and are expected to be detected simultaneously with the EOF. The 2D runtime represents the separation based on hydrophobicity in the first dimension, while the CZE-MS migration time represents the separation based on mobility.

Figure 2 A shows the base peak electropherogram (BPE) of the second dimension, together with zoomed extracted ion electropherograms (EIEs) of three different parts of the measurement. Each of the zoomed EIEs of the second dimension shows further separation, underscoring the orthogonality of the two dimensions. Figure S 2 clearly shows that the elution order from the one-dimensional nanoLC measurement is retained in the two-dimensional nanoLCxCZE-MS platform with only minimal peak broadening. We therefore assume negligible mixing in the storage capillary, consistent with findings of Bache et al. for RPLC gradient storage ^[28]^.

For a comprehensive evaluation of the 2D separation compared to the one-dimensional measurements, we evaluated the orthogonality based on the proteoforms identified by ProSightPD. Since we used a multisegment injection and a continuous MS acquisition, migration times of the second-dimensional measurements had to be calculated based on the 2D valve time program for injections to the second dimension. The calculated migration times across the two-dimensional measurement are shown in Figure 2 B and also highlight the high orthogonality. Taking the peak capacity of the nanoLC (60) and the peak capacity of the CZE (37), an overall peak capacity for the nanoLCxCZE-MS approach is calculated to be > 2200.

We evaluated the 2D nanoLCxCZE-MS method by comparing the number of proteoforms identified in the Caco-2 sample with the 1D nanoLC-MS and CZE-MS approaches. The measurements were performed by injecting samples with three different concentrations (an undiluted sample of ∼0.4 µg/µL, and a 1:5 and 1:10 dilution, corresponding to ∼4 µg, 0.8 µg, and 0.4 µg protein injected on the column) and analyzing the proteome by nanoLCxCZE-MS in the ∼ 35 min window stored from the first dimension (Figure 3 A) using the pseudo-comprehensive approach introduced in the experimental section of the supporting information.

**Figure 3:**
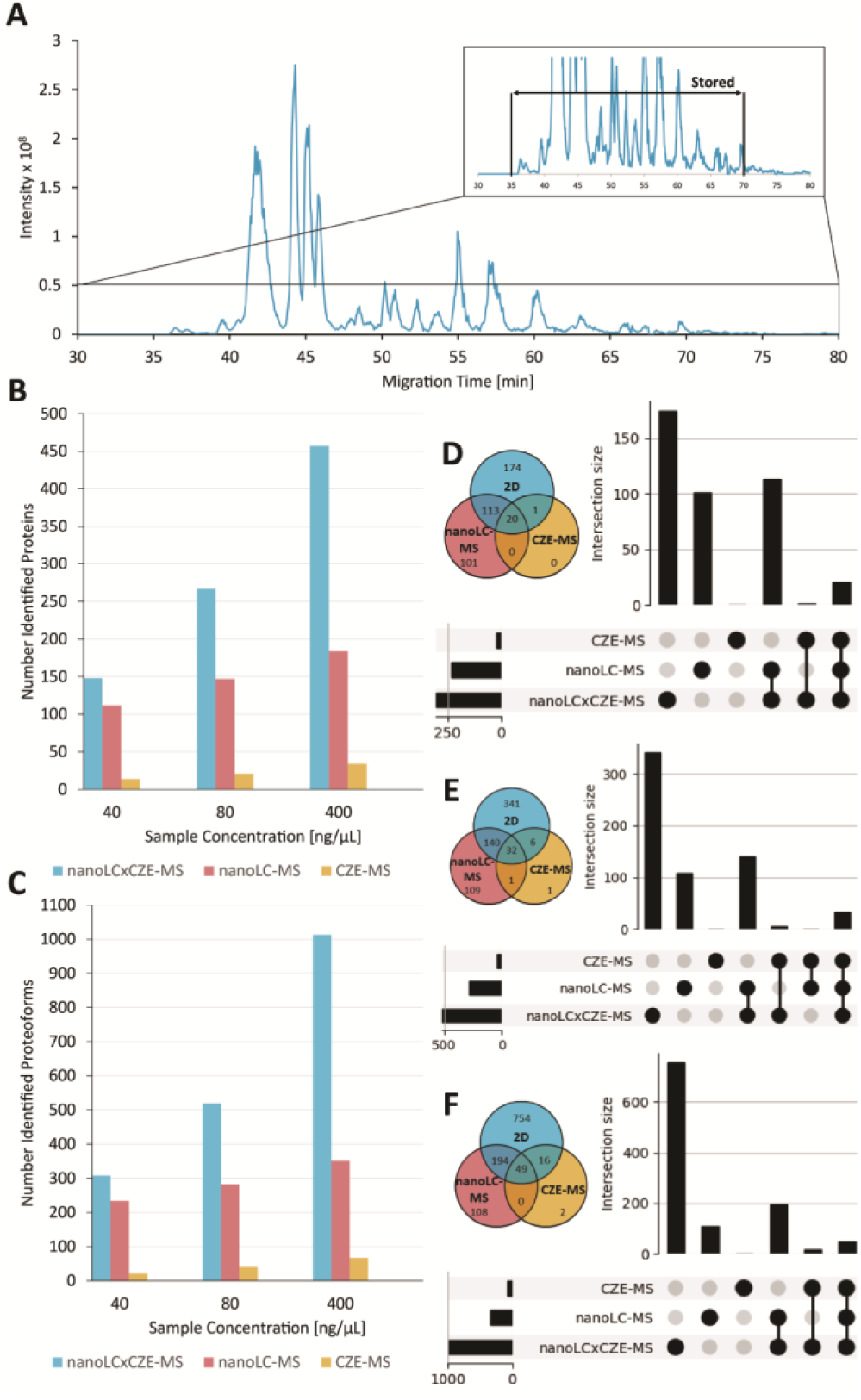
Sampled 1D region with the resulting number of proteins and proteoforms for different amounts of protein injected. A) 1D nanoLC-MS base peak chromatogram m/z 700-2000 with highlighted fraction characterized in detail using the 2D platform. B) Number of identified proteins for 2D nanoLCxCZE-MS, 1D nanoLC-MS, and 1D CZE-MS at three different sample concentrations. C) Number of identified proteoforms for 2D nanoLCxCZE-MS, 1D nanoLC-MS, and 1D CZE-MS at three different sample concentrations. D) Venn diagram and upset plot for the identified proteoforms at a sample concentration of ∼40 ng/µL. E) Venn diagram and upset plot for the identified proteoforms at a sample concentration of ∼80 ng/µL. F) Venn diagram and upset plot for the identified proteoforms at a sample concentration of ∼400 ng/µL. Intersection was determined using the Proforma entry from the Proteome Discoverer table. Proteoform identification parameters and workflow are described in the experimental section of the supporting information.

All three replicates of the nanoLCxCZE-MS data outperformed the 1D nanoLC-MS for the number of identified proteins (up to two times more using the nanoLCxCZE-MS platform, Figure 3 B) and the number of proteoforms (up to three times more using the nanoLCxCZE-MS platform, Figure 3 C). The average number of detected proteins and proteoforms per measurement, as well as the standard deviation of the triplicates, can be found in Figure S 3. The higher the amount of protein injected, the more the 2D platform outperformed the nanoLC-MS. This goes along with a strong increase in uniquely identified proteins and proteoforms by nanoLCxCZE-MS up to 754 uniquely identified proteoforms using the nanoLCxCZE-MS platform (Figure 3 D-F). 1D CZE-MS measurements identified only a low number of proteins (14-34 proteins depending on the sample concentration) and proteoforms (21-67 proteoforms depending on the sample concentration); almost all of them were also identified by the 2D measurement (Figure 3 D-F). We assume that the weaker performance of 1D CZE-MS is primarily due to its lower sample loading capabilities (a factor 160 lower amount of protein injected into the 1D CZE-MS) compared to nanoLC and nanoLCxCZE-MS. Thus, the online coupled nanoLC enables the analysis of low-concentration samples by CZE-MS.

Next, we performed triplicate comprehensive nanoLC×CZE–MS measurements of the undiluted sample (∼4 µg protein on column) to compare the number of identified proteins and proteoforms with the pseudo-comprehensive approach. The reduced pulsed volumes drastically increase the total measurement time, with ∼1470 min for the comprehensive vs. ∼670 min for the pseudo-comprehensive method. Although, applying the comprehensive nanoLCxCZE-MS method, we could increase the total number of detected proteins by ∼20 % to ∼550 and proteoforms by ∼ 16% to ∼1200 compared to the pseudo-comprehensive method (grouped processing of the triplicates Figure 4 A), considering the individual processed triplicates the differences between the two methods are not statistically significant (p=0.25 for proteins and p=0.19 for a two-sample t-test assuming equal variances and a 5% significance level (ttest2 in MATLAB R2024b). The average number of proteins and proteoforms identified by the individually processed measurements, as well as the standard deviation of the triplicates, are shown in the supporting information. (Figure S 4).

**Figure 4:**
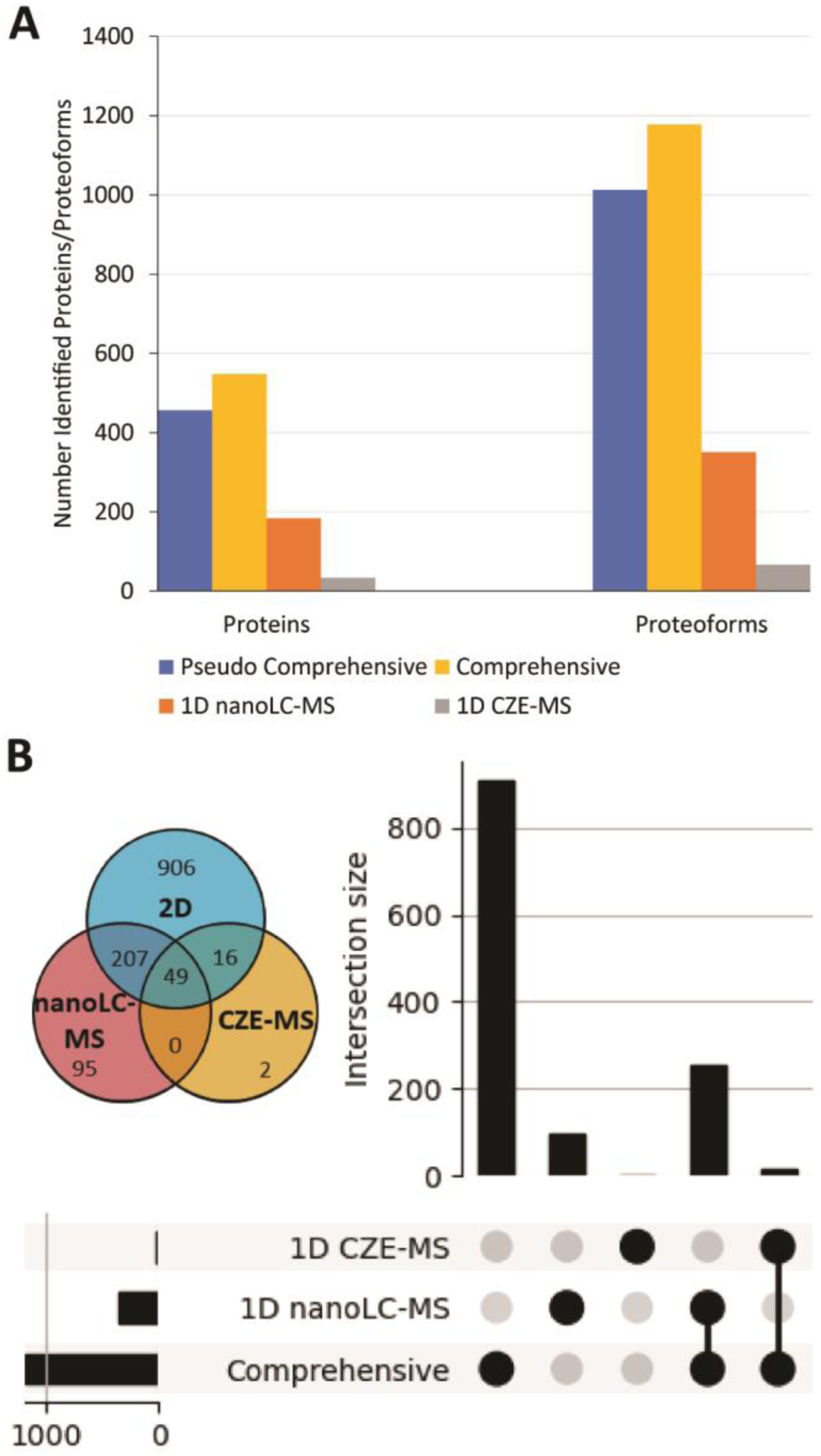
Number of proteins and proteoforms obtained for the measurement of the undiluted sample using 25 nL and 60 nL pulses. A) Number of identified proteins and proteoforms for 2D nanoLCxCZE-MS, 1D nanoLC-MS, and 1D CZE-MS (undiluted sample) grouped DB processing of the triplicates. The average number of detected proteins and proteoforms, as well as the standard deviation of the triplicates, can be found in Figure S4. B) Venn diagram and upset plot for the identified proteoforms of the undiluted sample for the comprehensive method. Intersection was determined using the Proforma entry from the Proteome Discoverer table. Proteoform identification parameters and workflow are described in the experimental section of the supporting information.

Due to the increased separation selectivity in the two-dimensional measurements, compared to the one-dimensional measurements, we identified ∼1200 proteoforms using our comprehensive two-dimensional approach. While the absolute number of identified proteins strongly depends on MS instrumentation and MS/MS settings, relative comparison between 1D and 2D measurements shows more clearly the benefit of our 2D platform: In the pseudo comprehensive, as well, as in the comprehensive approach, a 3-fold increase in detected proteoforms compared to 1D nanoLC-MS and a >15-fold increase compared to 1D CZE-MS were observed (Figure 4 A). Both in the pseudo-comprehensive data of all the different sample concentrations (Figure 3 D-F) and in the comprehensive data, approximately 100 proteoforms are only characterized in the 1D nanoLC-MS (Figure 3 D-F and Figure 4 B). To assess the complementarity, we analyzed the overlap between proteoforms identified by 1D nanoLC–MS and comprehensive nanoLCxCZE–MS. The comprehensive nanoLCxCZE–MS approach uniquely characterized 906 proteoforms (71% of the total identified proteoforms), demonstrating the substantially increased proteoform coverage enabled by the two-dimensional separation. Nevertheless, a subset of 95 proteoforms (7%) in the comprehensive measurement was exclusively identified in the 1D nanoLC–MS data (Figure 4 B). Of the 95 proteoforms uniquely characterized by 1D nanoLC-MS, 38 are less abundant and elute close to the edges or outside of the stored region, and could potentially be characterized by storing a different range from the first dimension. The remaining 57 proteoforms are unique for the 1D nanoLC measurement. Nevertheless, considering the 906 proteoforms uniquely characterized using the nanoLCxCZE-MS approach, our platform provides deep insight into the proteoform landscape.

### Mobility-based Proteoform and Proteoform Family Assignment

CZE separates proteoforms based on their electrophoretic mobility (i.e., charge-to-hydrodynamic size ratio), which can be calculated precisely in acidic separation conditions applied here, since only basic amino acids (and the N-terminus as well, as some PTMs) contribute to the protein charge at pH=2 ^[29,30]^. This includes charge-based separation of PTMs such as acetylation and phosphorylation, which often do not alter the hydrophobicity of the protein enough that these proteoforms can be separated by RPLC. Our platform, therefore, provides a tool for investigating proteoform families. For example, multiple acetylated histone H4 proteoforms are not separated by RPLC (Figure 5 A); however, they are baseline-separated by CZE (Figure 5 B). Note that the absolute intensity of the CZE-MS measurements is less compared to the nanoLC measurement due to the used sheath liquid interface, as shown in Figure S 6. Nevertheless, several additional proteoforms are detected at the MS1-level by nanoLCxCZE-MS, which were not detected in the 1D nanoLC-MS measurement. Moreover, nearly isobaric proteoforms containing either three methylations (Δm = 42.0470 Da, not changing the charge of the side chain) or one acetylation (Δm = 42.0106 Da, reducing the charge of the side chain) can be differentiated as shown in Figure 5 C. Although MS/MS information is missing, the nearly isobaric proteoforms can be tentatively identified by comparing the experimental with the calculated mobilities and assigned to the histone H4 proteoform family (Figure 5 D). This approach not only works for the discussed proteoforms but also for other peaks observed on the MS1-level, lacking sufficient MS/MS information for identification but most likely belonging to histone H4, since they form distinct groups regarding their retention and migration time and show distinct mass differences of 14 Da and 42 Da (Figure 5 E). If proteoforms tentatively identified on the MS1-level are found to be relevant, a subsequent targeted MS/MS analysis, e.g., in a pseudo-selective or heart-cut mode, could be carried out to obtain additional MS/MS information and characterize/validate the assigned proteoforms.

**Figure 5:**
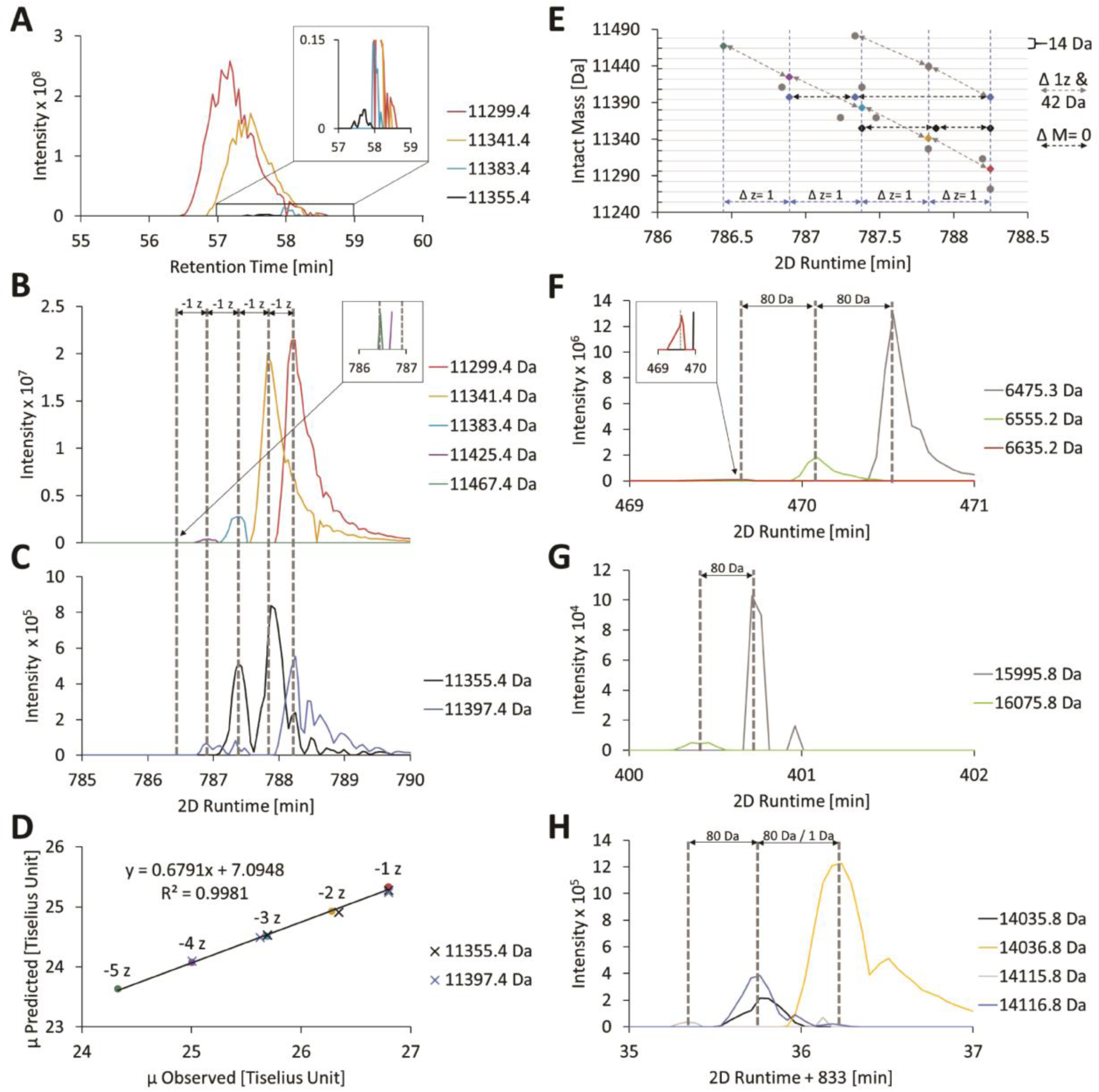
Separation of proteoforms in nanoLCxCZE-MS measurements. A) Extracted ion chromatograms of selected histone H4 proteoforms using 1D nanoLC-MS. B) Separation of selected histone H4 proteoforms assumed to vary by one charge due to acetylation based on MS1, in the second dimension of nanoLCxCZE-MS. C) Separation of isobaric proteoforms of histone H4 in the second dimension of the nanoLCxCZE-MS platform. Differences in migration times due to varying proteoform charges are indicated by dotted lines. D) Predicted vs. observed mobility of selected histone H4 proteoforms. Coloured dots predicted and observed mobility of the proteoforms from B, assumed to vary by one charge due to acetylation. Black and blue X predicted and observed mobility of the isobaric proteoforms from C. E) Intact mass vs. migration time of proteoforms attributed to histone H4, based on the MS1-level. Diamants represent proteoforms shown in D, black arrows indicate isobaric proteoforms, grey arrows indicate proteoforms varying in charge. F) Three proteoforms of keratin (P05787-1). Two of them were identified by ProSightPD, and one was postulated based on MS1 mass and migration time. G) Two proteoforms of Jupiter microtubule-associated homolog 1 (Q9UK76-1), one identified by ProSightPD processing, one postulated to carry an additional phosphorylation (postulated based on MS1 mass difference and migration time). H) Four proteoforms lacking sufficient MS/MS information for a detailed characterization. Mobility here proves valuable information to avoid wrong proteoform assignments.

In general, electrophoretic mobility can be used as an additional layer to validate identified proteoforms and to gain a more holistic picture of present proteoform families. Figure 5 F shows two proteoforms that were identified by ProSightPD as non-phosphorylated (6475.3 Da) and phosphorylated (6555.2 Da) keratin (P05787-1). The phosphorylated form with reduced charge and therefore reduced mobility migrates more slowly against the EOF than the non-phosphorylated proteoform (due to the reversed EOF, higher mobilities result in later migration times). In addition, a second phosphorylation is detected at the MS1-level (M = 6635.2 Da). Although the MS/MS information is missing for this proteoform, we can tentatively identify the PTM and assign it to a proteoform family, based on the combination of (i) similar elution time in the RPLC, (ii) matching mass difference on the MS1-level, and (iii) expected migration time difference. This approach also works for the 16 kDa Jupiter microtubule-associated homolog 1 (Q9UK76-1) shown in Figure 5 G.

Relative mobility not only allows for tentative identification of proteoforms but also for their validation. Figure 6H shows four proteoforms for which MS/MS information is missing. Based on the MS1-level, mass shifts indicate deamidation/ citrullination (1 Da mass shift) as well as phosphorylation (80 Da mass shift). Both are common PTMs, usually not well separated from their unmodified species in reversed-phase chromatography. Our method provides important additional information for validating PTMs due to the separation of mobility (charge to size ratio). While the 80 Da mass shift also shifts the migration time as expected for a phosphorylation, the mass shift of 1 Da does not fit the usually linked deamidation or citrullination, which would result in reduced mobility corresponding to earlier migration. This example clearly shows that the 1 Da mass difference has to be caused by another modification, e.g., on the sequence level. Thus, mobilities can be used to validate MS/MS-identified proteoforms, reducing the number of false positive proteoforms, which is a key problem in top-down proteoform analysis ^[29,31]^.

**Figure 6:**
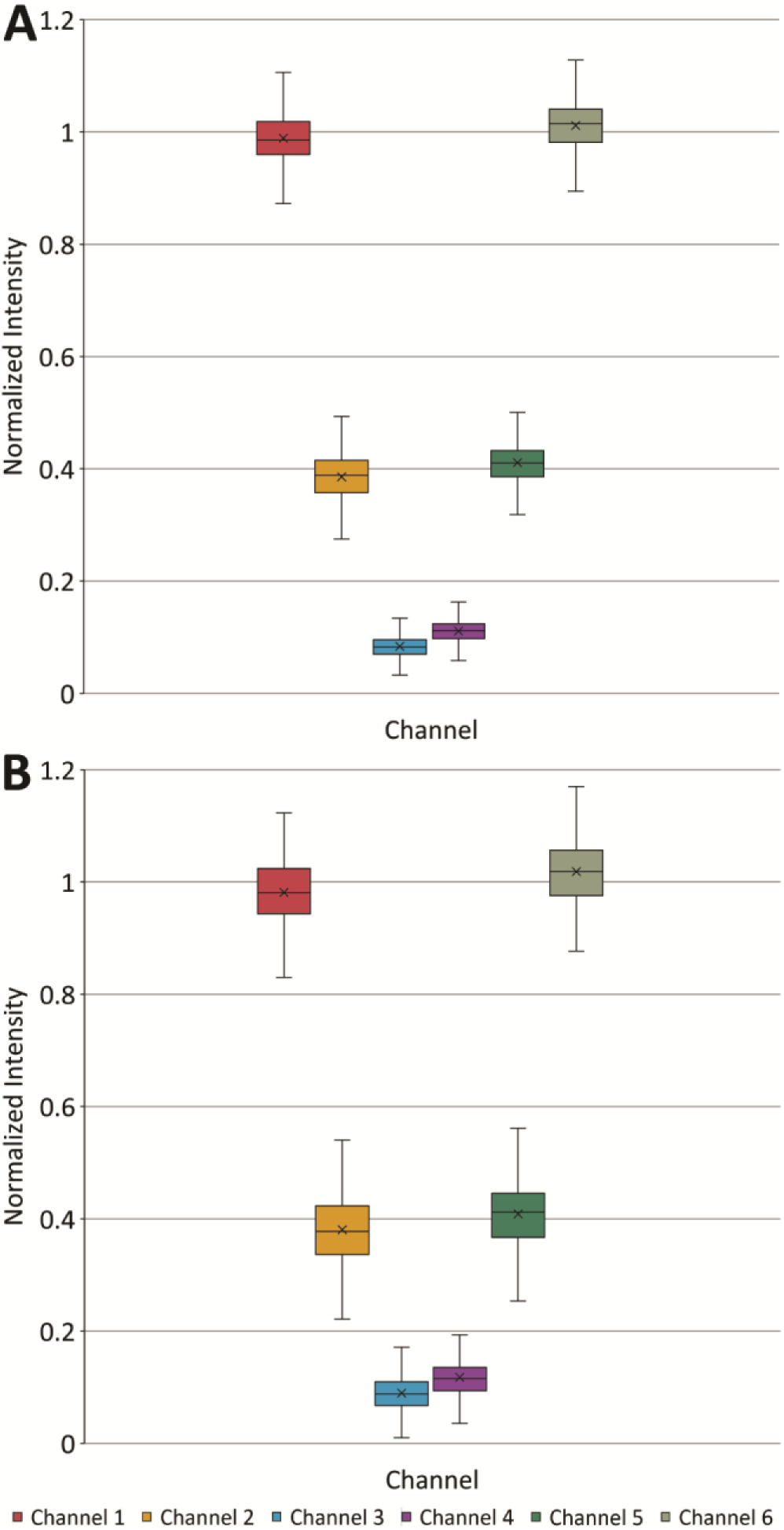
Quantification of E. coli proteoforms labelled with iodoTMT and mixed in defined ratios. The reporter ion intensities of several quantification scans originating from the same mass feature (within a 10-ppm mass tolerance) were matched to the via ProSightPD identified proteoforms. The reported intensities of each quantification scan were normalized to the average intensity of channel one and channel six of this quantification scan. The normalized intensity distributions are shown. The boxplot shows the 25th and 75th percentiles, with the whiskers indicating the 1.5 interquartile range. The black lines represent the median, and the black X represents the average value. A) Quantification using 1D nanoLC-MS, B) Quantification using pseudo-comprehensive nanoLCxCZE-MS.

### Quantitative Analysis of Cysteine-Directed Isobaric Labeled Proteoforms and Robustness

To demonstrate the feasibility of quantitation using our pseudo-comprehensive nanoLC-CZE-MS approach, an *Escherichia coli* cell lysate was divided into six aliquots, labeled with iodoTMTsixplex reagents for cysteine labeling, and recombined at a defined ratio as previously discussed^[32]^. This sample was analyzed by both nanoLCxCZE-MS and 1D nanoLC-MS. For each measurement, 2 µL of the sample was injected. The averaged detected ratio was 8:3:1:1:3:8, for both the 1D nanoLC-MS and the 2D nanoLCxCZE-MS measurements (Figure 6). We therefore consider both techniques equally suitable for iodoTMT-based quantitation.

In total, 1074 cuts were performed during the presented measurements. Of those 1074 cuts, only four had to be discarded due to air bubbles in the second dimension (air bubbles in a CZE separation are causing current issues and hindering proper electrophoretic separation). Note that if a cut was affected by an issue, only this cut had to be discarded, and the other cuts of the measurements could still be carried out and evaluated. Therefore, the total error rate of the presented platform accounts for less than 0.4%.

## Conclusion

In this work, we present a robust, flexible, and automated platform for reproducible online comprehensive nanoLCxCZE-MS TDP of biological samples. To the best of our knowledge, this is the first online nanoLCxCZE-MS platform storing the full nanoLC separation followed by subsequent time-decoupled TDP CZE-MS analysis. The elution order from the one-dimensional nanoLC measurement is retained in the two-dimensional nanoLCxCZE-MS platform for an extended period of time with only minimal peak broadening. By storing the first dimension in a capillary, our platform provides the ability to adapt both the first and the second dimension independently, e.g., regarding both the LC (e.g., gradient slope) and CZE separation (e.g., capillary length and coating), depending on the research question. This flexibility in the second dimension not only enables comprehensive nanoLCxCZE-MS analysis but also various other 2D methods like heart-cut and selective comprehensive analysis, while still being flexible regarding the second-dimension modulation time. In addition, the high flexibility and modular structure of our platform allow us to easily switch between 1D nanoLC and nanoLCxCZE-MS.

Pseudo-comprehensive sampling substantially reduces analysis time while preserving most of the information. The use of iodoTMT labeled samples enables quantitative proteoform comparisons, also when performing pseudo-comprehensive nanoLCxCZE-MS analysis. The platform is automated, requires no manual sample handling or fraction collection, and can perform the analysis unsupervised with a low error rate of less than 0.4%.

Applying the here introduced novel nanoLCxCZE-MS platform to biological samples, we demonstrate high orthogonality between the dimensions (hydrophobicity and electrophoretic mobility). This results in a peak capacity of 2200 and a strongly improved separation of proteoforms belonging to the same proteoform family. Due to the improved separation, the risk of co-isolation during MS/MS, e.g., during isobaric label-based quantification, can be minimized. The number of proteoforms identified by data-dependent MS/MS experiments is increased by >3-fold compared to 1D nanoLC-MS and >15-fold compared to 1D CZE-MS. Since mobilities in the second dimension can be calculated accurately, it is possible to attribute protein masses to certain proteoform families if MS/MS information is not sufficient to identify the proteoform. This is shown exemplarily for the histone H4 family, where the proteoforms can be clearly attributed based on their mobility and their intact mass. Furthermore, mobilities can be used to validate MS/MS-identified proteoforms, reducing the number of false positive proteoforms. Compared to 1D RPLC-MS, our nanoLCxCZE-MS method provides additional power for both the assignment and detection of false assignments of proteoforms and proteoform families.

Overall, the presented platform is expected to be of great value for targeted and untargeted TDP, especially for applications where sensitive characterization of proteoforms and investigation of proteoform families in complex biological samples is aimed for. We expect the platform to further benefit from improved protein characterization capabilities, e.g., advanced MS instrumentation, also including ion mobility technology. In addition, improved algorithms for data-dependent MS/MS precursor selection will boost proteoform identification and characterization, since, e.g., for each cut, different precursors belonging to different proteoforms could be selected for fragmentation, and already fragmented proteoforms could be excluded from further fragmentation. With these improvements, the presented nanoLCxCZE-MS platform will be even more valuable compared to existing 1D workflows.

## Supporting Information

The experimental section, data analysis description, and additional supporting figures can be found in the supporting information file. The mass spectrometry proteomics data have been deposited to the ProteomeXchange Consortium via the PRIDE ^[33]^ partner repository with the dataset identifier PXD076819.

## Author Contributions

TW and CN conceived the study design. PTK and AT provided the samples. All experiments, as well as the preparation of figures and the initial manuscript was performed by TW. Data processing and evaluation by PTK and TW. All authors contributed to the discussions as well as the editing and formatting of the manuscript.

## Acknowledgments

This project was funded by the Deutsche Forschungsgemeinschaft (DFG, German Research Foundation; 465662382).

## Notes

### Competing Interest Statement

The authors have declared no competing interest.

