## Supporting_Information for "Comprehensive online two-dimensional nanoLCxCZE-MS for deep top-down proteomics"

### Experimental Section

#### Chemicals and Materials

Ultrapure water ($H_2O$, 18 MΩ*cm at 25 °C) was prepared using an SG Ultra Clear UV (Siemens Water Technologies, USA). Acetonitrile (LC-MS grade), 2-propanol (LC-MS grade), acetic acid (HAc, 100%), and formic acid (FA,≥98%) were purchased from Carl Roth (Karlsruhe, Germany). Poly(diallyldimethylammonium chloride) (PDADMAC, Mw 400,000-500,000 g/mol), Poly(styrene sulfonate) (PSS, Mw 70,000 g/mol), 2-[4-(2-hydroxyethyl)piperazin-1-yl]ethane sulfonic acid, hydrofluoric acid (40% (v/v)), sodium hydroxide, hydrochloric acid (37%), Myoglobin from equine skeletal muscle (purity ≥ 95%), Ribonuclease A from bovine pancreas (purity ≥ 60%), and Lysozyme from chicken egg white (purity ≥ 90%) were purchased from Sigma-Aldrich. Fused silica pulled tip ESI Emitters with 10 µm Orifice ID were purchased from CoAnn Technologies Inc. (Richland, US). Bare fused silica (FS) capillaries with 50 µm inner diameter (ID) and 375 µm outer diameter (OD) were acquired from Polymicro Technologies (Phoenix, AZ, USA). Glass emitters with 1 mm OD and an orifice ID of 30 µm were obtained from BioMedical Instruments (Zoellnitz, Germany). Methyl-Sil deactivated FS capillaries with 50 µm ID and 360 µm OD were purchased from CS-Chromatographie Service GmbH (Langerwehe, Germany).

**Samples**

For 1D and 2D measurements, proteins from human Caucasian colon adenocarcinoma cells (CaCo-2) were used. The cell lysate was purified by solid-phase extraction; thus, primarily proteoforms smaller than approximately 20 kDa were enriched during sample preparation[1]. Ribonuclease A, Lysozyme, and Myoglobin were used to evaluate and monitor the CZE-MS dimension. Protein stock solutions (4 mg/mL in $H_2O$) were mixed and diluted to 200 µg/mL per protein using 2 M acetic acid.

a) Cell Cultivation and CaCo-2 Sample Preparation

RPMI-1640 medium, fetal bovine serum albumin, and TrypLE™ Express Enzyme were purchased from Thermo Fisher Scientific (Bremen, Germany). cOmplete EDTA-free protease inhibitor cocktail was purchased from Roche (Penzberg, Germany). All other chemicals and Caucasian colon adenocarcinoma (CaCo-2) cells (ATCC Number: HTB-37™) were purchased from Sigma-Aldrich (Steinheim, Germany). Deionized water (18.2 MΩ/cm−1) was prepared using an arium611 VF system (Sartorius, Göttingen, Germany).

CaCo-2 cells were maintained as per European Collection of Cell Cultures (ECACC) recommendations and as previously described[2]. The cells were grown in RPMI-1640 medium (25 mM HEPES, 2 mM L-glutamine, 0.013 mM phenol red) supplemented with 10% (v/v) fetal bovine serum, and 1% (v/v) penicillin (10,000 U/mL)/streptomycin (10,000 µg/mL) at 37 °C with 5% CO2. The cells were passaged between 90-100% confluence and detached using TrypLE™ Express Enzyme. Cells were washed three times with PBS buffer (centrifugation at 200×g, 5 min at 25 °C) before harvesting. Cell pellets were stored at −80 °C until cell lysis. Enrichment of the low-molecular-weight fraction of the proteome was performed by solid-phase extraction as previously described [2,3]. In brief, ca. $2\times10^7$ CaCo-2 cells were lysed in freshly prepared unbuffered 8 M guanidinium chloride (GndHCl; to prepare the stock solution, an appropriate amount of GndHCl was dissolved in MilliQ water and heated for 30 min to 35 °C) supplemented with cOmplete protease inhibitor by freeze-thaw cycling (10× 1 min −80 °C S3 ethanol bath, 1 min sonification). The sample was acidified with 1 ml 5% formic acid (FA), and centrifuged (21,100 g, 15 min). The supernatant was transferred to a preconditioned SepPak (3cc, C18, washed 2× 3 ml with 100% acetonitrile and 2× 3 ml 5% FA) and washed twice with 3 ml 5% FA. Proteoforms were eluted with 300 µl 70% ACN (0.1% TFA) and 100% ACN. The eluate was dried to complete dryness by lyophilization and resuspended in 3% ACN, 0.1% FA before analysis.

The protein concentration of the resuspended sample was determined using the Pierce BCA Protein Assay Kit (Thermo Scientific).

b) Cell Cultivation and Lysis of Escherichia coli (E.coli) Cells

*Escherichia coli* K-12 strain MG1655 was cultured in M9 minimal medium supplemented with 15 mM glucose as previously described [4]. Bacterial cultures were started at an initial optical density at 600 nm (OD600) of 0.1 and grown at 37 °C with continuous shaking. Cells were collected once the culture reached an OD600 of approximately 1 by centrifugation (3 min, 3,000 rcf, room temperature). The resulting cell pellet was rinsed with ultrapure water and stored at −80 °C until further processing.

c) Escherichia coli Sample Preparation and Labeling

*E. coli* cells were resuspended in lysis buffer (6.4 M guanidinium hydrochloride, 200 mM triethylammonium bicarbonate (pH 8.5), 1× cOmplete protease inhibitor (Roche, Basel, Switzerland)), and lysed by ultrasonication on ice. After centrifugation (21,100 rcf, 20 min,

4 °C), the protein concentration of the supernatant was determined using the Pierce BCA Protein Assay Kit (Thermo Scientific).

Cysteine-directed iodoTMT labeling was performed as previously described [5]. Briefly, 160 µg protein was transferred into a reaction tube for each TMT channel. The volume was filled up to 80 µL with lysis buffer. Proteoform reduction was performed by adding 1 µL 200 mM Tris(2-carboxyethyl)-phosphine (TCEP) and incubation for 60 min at 50 °C and 800 rpm on a shaker. The iodoTMTsixplex reagents (Thermo Scientific) were resuspended in 10 µL methanol and added to the samples, followed by incubation for 50 min at 37 °C and 800 rpm in the dark. The reaction was quenched by adding 4 µL of 200 mM DTT and incubation at room temperature for 15 min. Then, the samples were combined and purified by methanol-chloroform water precipitation. The protein pellet was resuspended in 97% water, 3% ACN+ 0.1%TFA before injection.

**Nano-Liquid Chromatography (nanoLC)-MS**

For nanoLC separations, an UltiMate™ 3000 RSLCnano system (Thermo Fisher, Germering, Germany) was used. Separation was carried out using a customized ReproSil Gold 300 C4, 3 µm, 450 mm × 75 µm column equipped with a ReproSil Gold 300 C4, 3 µm, 80 mm × 75 µm precolumn (Dr. Albin Maisch High Performance LC GmbH, Ammerbuch, Germany). The analytical column and the precolumn were in-house slurry-packed according to the FlashPack protocol [6]. Eluent A was composed of 100% $H_2O$ + 0.1% FA, and eluent B of 80% acetonitrile, 20% $H_2O$ + 0.1% FA. Large volume injection was performed using a ReproSil Gold 300 C4, 5 µm, 10 mm × 0.3 mm trap guard column (Dr. Albin Maisch High Performance LC GmbH). 10 µL sample was loaded on the trap column with 5% Eluent B at a flow rate of 20 µL/min for 5 min. Separation was carried out with a linear gradient of 55 min from 10% to 95% Eluent B at a nanoLC flow rate of 100 nL/min. The separation column was flushed with 95% Eluent B for 15 min before re-equilibration at 10% Eluent B for 64 min. For 1D nanoLC-MS experiments, the nanoLC was coupled to an Orbitrap Fusion Lumos Tribrid mass spectrometer (Thermo Fisher Scientific) using a home-built interface based on the UWPR nanospray source design from the University of Washington (https://proteomicsresource.washington.edu/protocols05/nsisource.php) and a pulled-tip fused silica ESI emitter with a 10 µm Orifice ID (CoAnn Technologies Inc., Richland, US). NanoESI spray voltage was set to 1.8 kV. All data were acquired in the intact protein, low-pressure, and positive MS mode. For MS1, a mass range of 500–2000 *m/z* and a resolution of 120,000 were used. The normalized automatic gain control target was set to 200%, and the maximum injection time to 246 ms with four microscans. For MS/MS, a scan range of =150–2000 *m/z* and a resolution of 60,000 were used. The normalized automatic gain control target was set to 1000%, and the maximum injection time to 250 ms with four microscans. The ions were isolated using a 3 *m/z* isolation window in the quadrupole and a 650-2000 m/z and 4-50 charges precursor filter. Precursors were fragmented using electron transfer higher energy collisional dissociation (EThcD). Electron transfer dissociation (ETD) reaction time was 10 ms (ETD reagent target of $6E^5$ with maximal ETD reagent injection time of 200 ms), followed by 23% normalized higher energy collisional dissociation (HCD) collision energy. For IodoTMT quantification, in addition to the EThcD scan, a quantification scan was performed at 80% normalized higher energy collisional dissociation (HCD) collision energy, resolution 30,000, and a MS/MS scan range of 120-140 *m/z*. The normalized automatic gain control target for the quantification scan was set to 1000%, and the maximum injection time to 54 ms with one microscan.

**Capillary Zone Electrophoresis (CZE)-MS**

CZE-MS was performed as previously described[2] with slight modifications. For CZE-MS, a G1600 HP 3D CE instrument (Agilent Technologies, Waldbronn, Germany) and coated 50 µm-ID capillaries were used. The tip of the separation capillary was etched with hydrofluoric acid to an outer diameter of ~ 80–100 µm to sufficiently penetrate the emitter. The capillaries were coated using a five-layer PDADMAC-PSS successive multiple ionic polymer layer (SMIL) according to the protocol of Dhellemmes et al.[7]. To stabilize the coating, −15 kV was initially applied for 30 min. Prior to analysis, the capillary was flushed with background electrolyte (BGE, 2 M HAc).

QC and 2D measurements were performed using a separation voltage of −15 kV, a 33 cm connection capillary, a 35 cm separation capillary, and the 2D valve. 1D CZE-MS measurements were performed at −20 kV using a 65 cm capillary without the 2D valve. 1D CZE-MS injection was performed by applying 50 mbar for 48 seconds.

The CE was coupled online to an Orbitrap Fusion Lumos Tribrid mass spectrometer using the nanoCEasy interface[8] equipped with 30 µm orifice ID emitters. Isopropanol: $H_2O$ (50:50, v/v) + 0.5% FA was used as a sheath liquid. The spray voltage for the nanoCEasy interface was set to +2 kV for all 1D and 2D CZE-MS measurements. All other MS parameters were the same as for the nanoLC-MS measurements.

**(Selective) Comprehensive nanoLCxCZE-MS platform**

The nanoLC separation column was connected to a Methyl-Sil deactivated FS storage capillary with 50µm ID via the UltiMate 3000 RSLC column-switching valve. The other end of the storage capillary was connected to an 8-port 4-position 20 nL PEEK valve (2D-valve, VICI AG International, Schenkon, Switzerland). The adjacent ports of the 2D valve were connected to two 50 µm ID SMIL-coated capillaries for electrophoretic separation. The SMIL connection capillary connecting the CE instrument to the 2D valve was 33 cm long, while the separation capillary from the 2D valve to the mass spectrometer was 35 cm long. Chromatographic separation in the first dimension was carried out as described above. CZE separation was carried out at −15 kV. Switching of the 2D valve was triggered based on the chosen modulation time of the second dimension by an in-house written script running on a Raspberry Pi 4 model B (Raspberry Pi Ltd, Cambridge, UK). The unused ports were flushed with $H_2O$ using 50 µm ID capillaries and a NE-1002XES microfluidics syringe pump (Adaptas by SIS, Massachusetts, US) equipped with a 5 mL syringe (Trajan Scientific and Medical, Victoria, Australia). All MS parameters were the same as for the CZE-MS measurements. Due to the Xcalibur method limit (999 min), the MS measurement had to be split into two for the comprehensive analysis using 25nL pulses. Chromeleon limits the total duration of a 2D measurement to 1500 min.

**Data Analysis**

Manual MS data analysis was carried out in Freestyle 1.8 (Thermo Fisher Scientific) using the XTRACT deconvolution algorithm. Data were deconvoluted using a charge range of 5-50, the FreeStyle Protein isotope table, and a minimum number of detected charges of 3. If necessary, individual extracted ion chromatograms and electropherograms were smoothed by applying a Gaussian smoothing algorithm using a smoothing level of 5 in FreeStyle 1.8.

Proteoform identification was performed with ProSightPD (Version 4.2, Proteinaceous, Inc., Evanston) within Proteome Discoverer (Version 3.0.0.757, Thermo Scientific, Germany) using a 1% false discovery rate. Deconvolution of the raw data was performed using the High/High cRAWler (Xtract), and proteoform identification was performed against a human protein database (only reviewed proteins downloaded from UniProt as an XML file, including all known modifications, taxon-ID 9606, release 2023_01) using the Annotated and Subsequence Proteoform Search nodes (precursor and fragment mass tolerance: 10 ppm).

For quantitative evaluation, the raw files were converted to the mzML format using msConvert [9], with peak picking enabled. Deconvolution and mass feature TMT ratio extraction were performed with FLASHDeconv in FLASHApp (version 0.9.15) [10–12]. The quantitative information from the FLASHDeconv output was matched to the cysteine-containing proteoforms identified using ProSightPD based on their reported mass (within a 10-ppm tolerance). The reported intensities of each quantification scan were normalized to the average intensity of channel one and channel six of this quantification scan.

#### Method Optimization

Notably, by storing the first-dimension separation in a capillary, the analysis time in the second dimension can be chosen independently. Nevertheless, to achieve the best efficiency (i.e., the highest possible sample throughput in a given time), several aspects need to be considered.

a) nanoLC Runtime

The nanoLC runtime is primarily determined by the column length and the applied gradient. For our instrument configuration, a linear gradient that elutes proteins over a 35 min window is used; however, this can be selected according to the application.

b) CZE-MS Runtime

Applying a given voltage, the separation time of CZE depends on the length of the capillary and overall apparent mobilities (resulting mobility of an analyte, when considering the mobility of the analyte and the mobility of the (here counter-directed) electroosmotic flow (EOF)). For the commercial CE instrument, a 33 cm capillary is required to connect the CE instrument to the 2D valve, and a 35 cm separation capillary to connect the 2D valve to the MS. The overall runtime of the CZE-MS was ~16 min, applying a separation voltage of 15kV and a highly efficient cationic 5-layer SMIL coating (PDADMAC: PSS), leading to a relatively high EOF towards the MS. The use of the high EOF in our 2D approach greatly simplifies the CZE protocol, since no flushing steps are required due to the permanent exchange of the BGE in the capillary by the EOF.

c) Reduction of Void Times in the CZE-MS by Multisegment Injection

The observed migration times of our analytes ranged from ~ 5 min to ~16 min, resulting in a window of ~11 min where analytes are detected (Figure 2 B). Therefore, we applied multisegment injection to further reduce cycle time and increase throughput in the second dimension. The concept was first introduced by Kuehnbaum et al. for metabolomics using CZE-MS [13] and later also used for TDP by Zhao et al. [14]. In the classical multisegment approach, multiple samples are injected hydrodynamically with spacers in between sample segments. Here, we implemented multisegment injection using a continuous CZE method since the sample loop becomes part of the CZE dimension after valve switching, and a high-EOF coating is used. Every 11 minutes, the separation voltage in the second dimension was reduced to 0 V, and the next sample was injected by switching the 2D valve. BGE spacer segments were generated by the high EOF system automatically in between injections. By applying multisegment injection, we reduced the cycle time in the second dimension from ~16 min to 11.5 min, thus reducing the overall 2D runtime by ~30%. Therefore, this approach was applied to all the measurements.

d) Sampling Frequency From the nanoLC Separation

The mean peak width of the most intense first-dimensional nanoLC peaks was 1.4 ± 0.6 min (baseline, n=3 samples with 19 peaks each, Figure S 2 A), corresponding to a peak volume of ~140 nL. Thus, the 20 nL transfer volume of the CZE will result in several CZE-MS measurements from the same LC-peak. To fully characterize a sample, it is sufficient to transfer each peak of the nanoLC once into the CZE-MS ("pseudo comprehensive"), rather than transfer each volume from the first dimension to the second dimension ("full comprehensive"). Pulsing 60 nL to the storage capillary, from which only 20 nL can be transferred to the second dimension, still sampled each peak 3 ± 1 times (n = 3 samples with 19 peaks each), as shown in Figure S 2 C. A close-to full comprehensive experiment (25 nL pulsed volume from which 20 nL are transferred to the second dimension, in this publication referred to as "comprehensive" for simplicity's sake) just increases the number of repeated detections of one protein to 8 ±3 (n=3 samples with 19 peaks each, Figure S 2 D+E). Thus, most experiments have been performed in the pseudo-comprehensive mode with a pulse volume of 60 nL, saving about 60% in time compared to the comprehensive mode. The sampling frequency (i.e., the pulse volume) from the nanoLC separation can be selected over a wide range with high reproducibility. The average pulse volume over the twelve discussed measurements performed at 60 nL/cut was 60±4 nL/min (determined as average pulse/cut over each of the 2D measurements). Average pulse volume over the three measurements performed at 25 nL/cut was 25±1 nL/min (determined as average pulse/cut over each of the 2D measurements). An exemplary calculation of the pulse volume is shown in the supporting information.

**Calculation of Peak Capacity**

1D Peak capacity was calculated based on the average FWHM of the 19 most intense one-dimensional peaks shown in Figure S 2 A and the 1D retention time window (~33 min) spanned by those 19 proteoforms. 2D Peak capacity was calculated based on the average FWHM of the 19 proteoforms in the second dimension, Figure S 2 D+E, and 11 min per cut.

**Calculation of Pulsed Volume**

To determine the experimental pulsed volume, the volume between the peak maxima of two proteoforms was divided by the number of cuts between the most intense cuts of those proteoforms in the second dimension. E.g., proteoform A elutes at 39.45 min, and proteoform B elutes at 66.58 min in the first dimension. This accounts for a volume of 2700 nL. The most intense cut of proteoform A is detected at ~75 min, and the most intense cut of proteoform B is detected at ~588 min in the second dimension. The time difference between the most intense cuts of those proteoforms is 513 min, which accounts for 45 cuts. 2700 nL/45 cuts = 60 nL/cut.

### Supplementary Notes

#### Operation Modes of the Platform.

Besides 1D nanoLC-MS and 1D CZE-MS, the platform can be used for various 2D applications, ranging from heart cut (fast but limited information about the sample) to comprehensive (time-consuming but deep information about the sample). A selective comprehensive mode (Figure S 1 B) is useful for a targeted approach to characterize only a certain LC fraction and, thus, save time. This mode can also be used to sample different non-consecutive peaks of the first dimension (Figure S 1 C). We applied this mode, e.g., for the characterization of different Histones (data not shown). The comprehensive approach (Figure S 1 D) is the method of choice for untargeted deep proteome analysis with minimal prior knowledge about the sample.

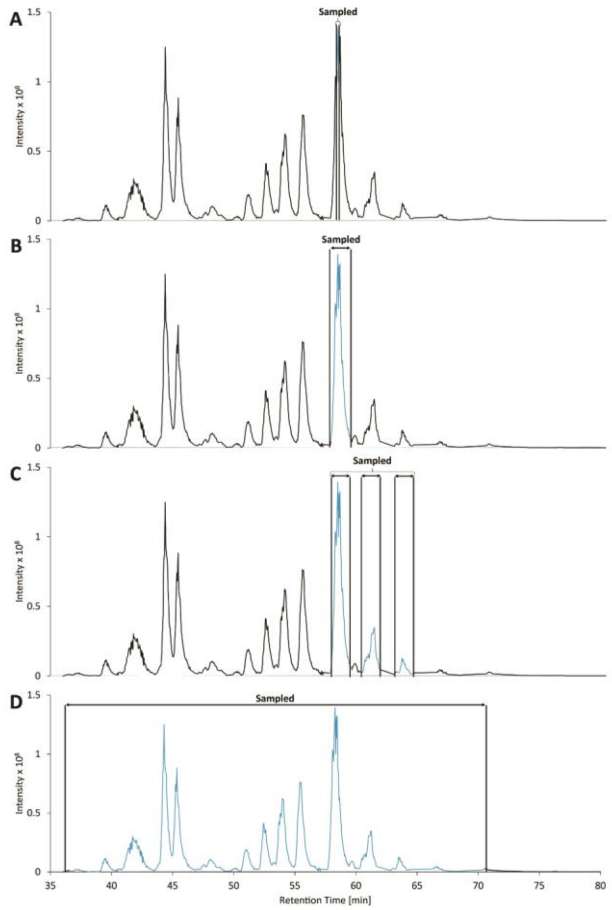

*Figure S 1: Deconvoluted base peak chromatograms for possible 2D modes of the nanoLCxCZE-MS platform. A) Heart-cut analysis of one Peak. B) Analysis of one Peak in the selective comprehensive mode. C) Analysis of three different parts of the chromatogram using the selective comprehensive mode. D) Comprehensive analysis of ~35 min (~3.5 µL) from the first dimension.*

**Comparison of 1D and 2D Measurements Based on Selected EIE**

Based on the base peak chromatogram (BPC) after deconvolution of the 1D nanoLC-MS measurement, the most intense mass of the BPC-peaks was noted. Next, extracted ion chromatograms (EICs) and extracted ion electropherograms (EIEs) with 10 ppm tolerance were generated for those masses Figure S 2. The EIC of the nanoLC represents the elution order of the proteins, Figure S 2 A. The different underlying separation mechanisms can be highlighted by the different migration orders in 1D CZE-MS (Figure S 2 B). In addition, the EIEs of the two-dimensional measurements (Figure S 2 C-E) show that we retain chromatographic separation during storage and cut each BPC peak multiple times in the case of the 60 nL/cut measurements (Figure S 2 C). The EIEs also highlight that the peaks of the first dimension are cut more often if we reduce the pulsed volume to 25 nL/cut (Figure S 2 D+E). Please note that due to software limitations, the mass spectrometric detection of the 2D measurement with 25 nL/cut pulsed volume had to be split into two parts (Figure S 2 D+E).

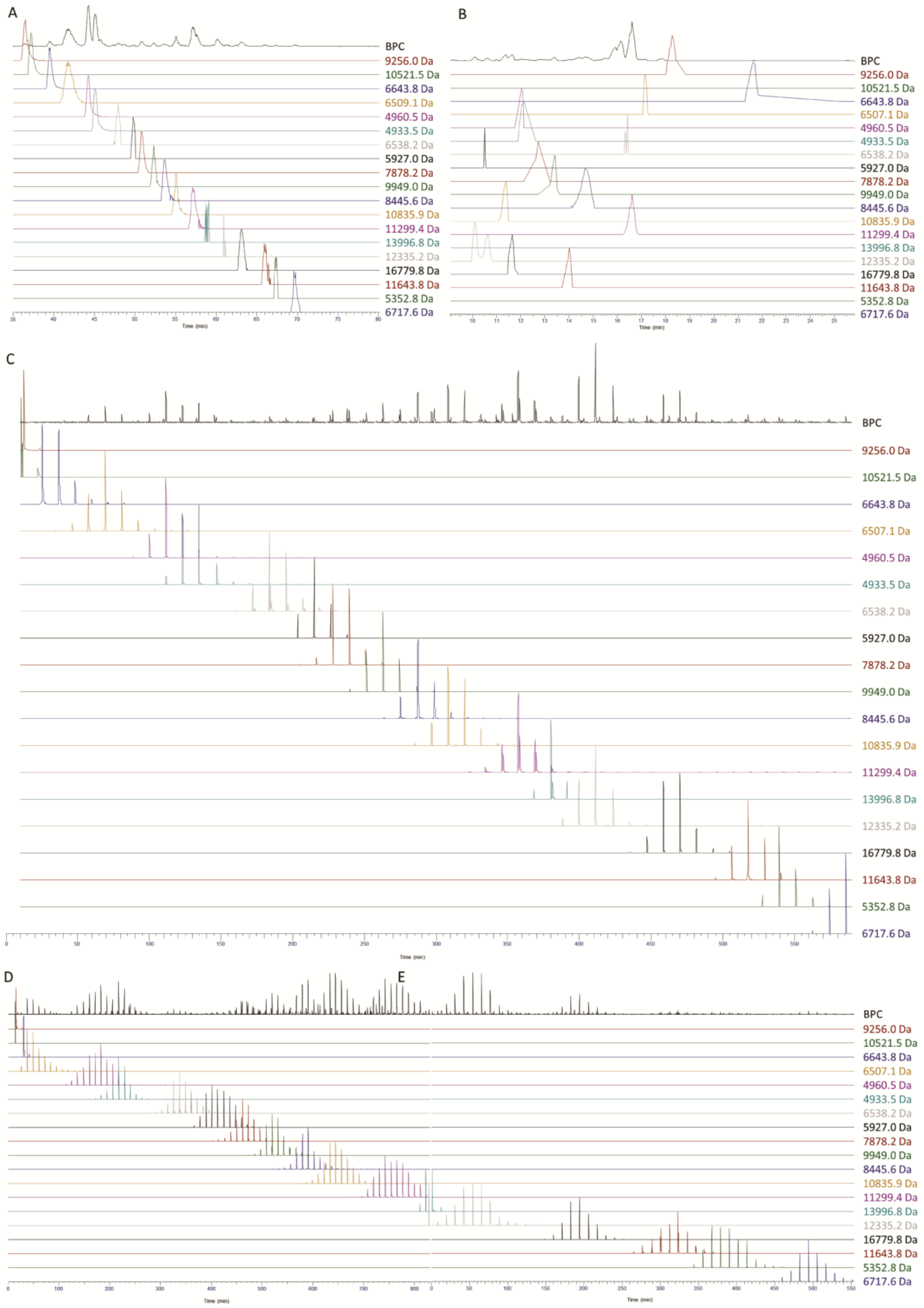

*Figure S 2: BPC/ BPE and selected EICs and EIEs of one-dimensional and two-dimensional measurements after deconvolution. A) 1D nanoLC-MS, B) 1D CZE-MS, C) 2D nanoLCxCZE-MS 60 nL pulsed volume/cut, and D) 2D nanoLCxCZE-MS 25 nL pulsed volume/cut part 1, E) 2D nanoLCxCZE-MS 25 nL pulsed volume/cut part 2.*

#### ID of Proteins and Proteoforms for Individual Measurements

Figure S 3 and Figure S 4 provide additional information to the figures shown in the main manuscript. While for the figures in the main manuscript the triplicate measurement was processed as a group, for the figures here, each measurement was processed individually. Bars show the average number of proteins and proteoforms, while the error bars represent the standard deviation of the triplicate measurements.

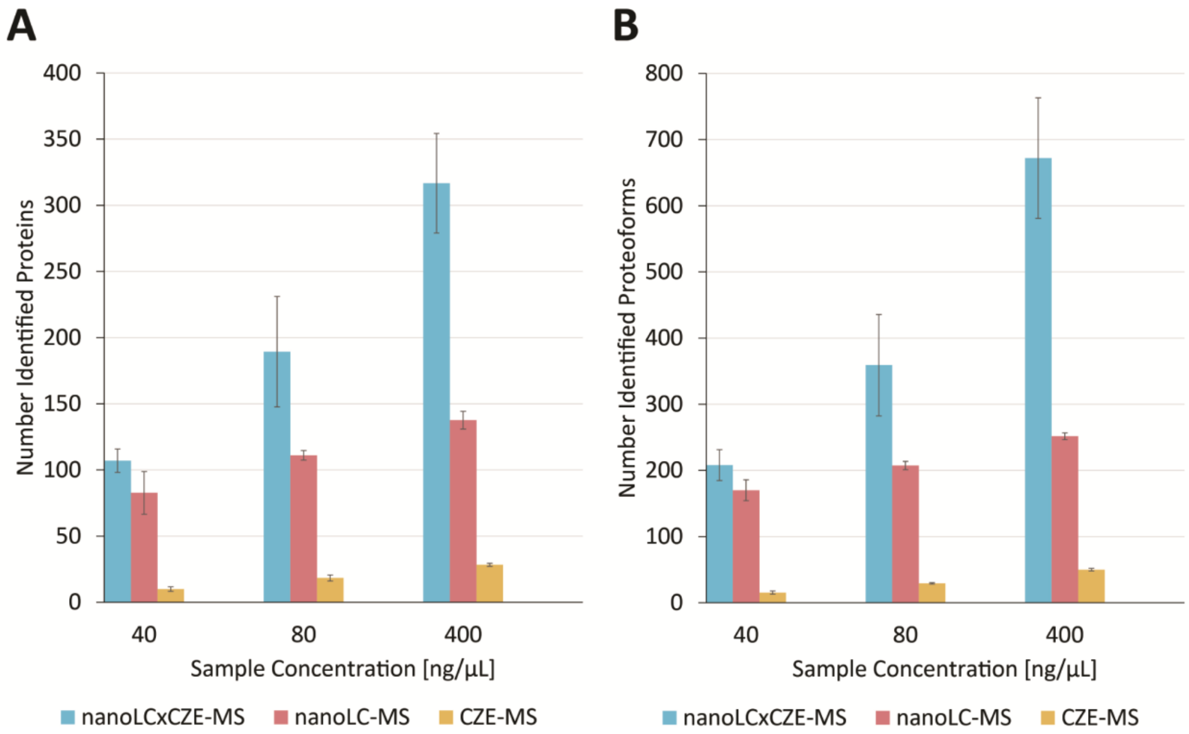

*Figure S 3: Average number of identified proteins and proteoforms with error bars (± standard deviation) for the 2D measurement using 60 nL/cut pulsed volume at different sample concentrations (~40 ng/µL, ~80 ng/µL, and ~400 ng/µL). A) Proteins, B) Proteoforms. Proteoform identification parameters and workflow are described in the experimental section.*

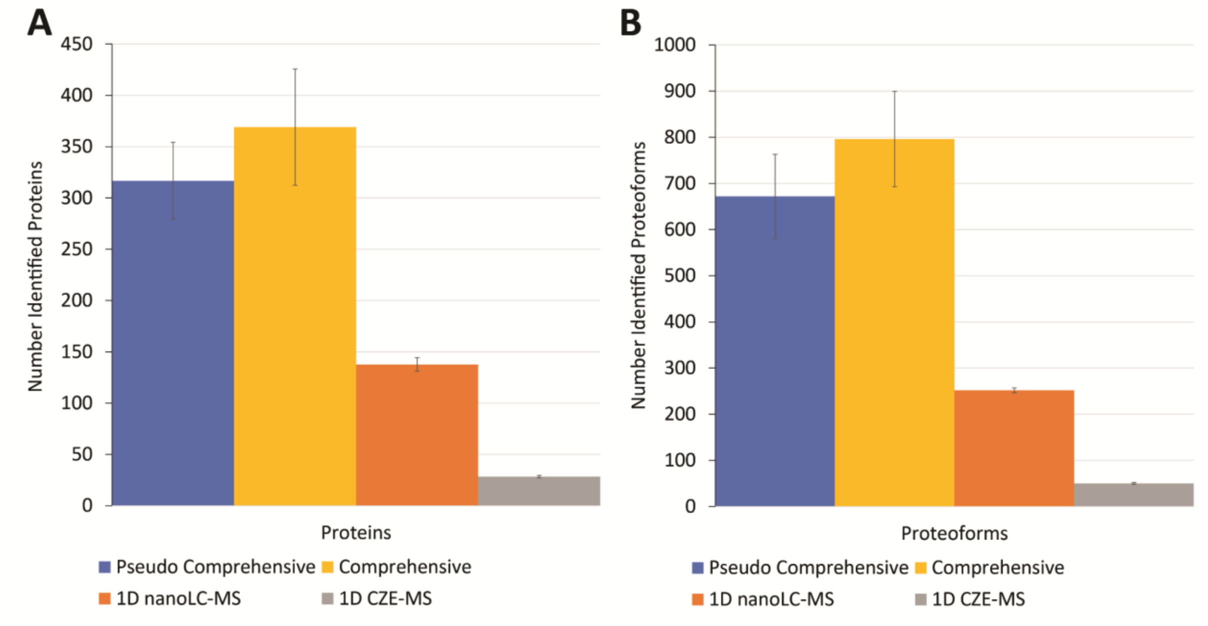

*Figure S 4: Average number of identified proteins and proteoforms with error bars (± standard deviation) using the undiluted sample. A) Proteins, B) Proteoforms. Proteoform identification parameters and workflow are described in the experimental section.*

#### Additional Information Figure 6 (Proteoforms Histone H4)

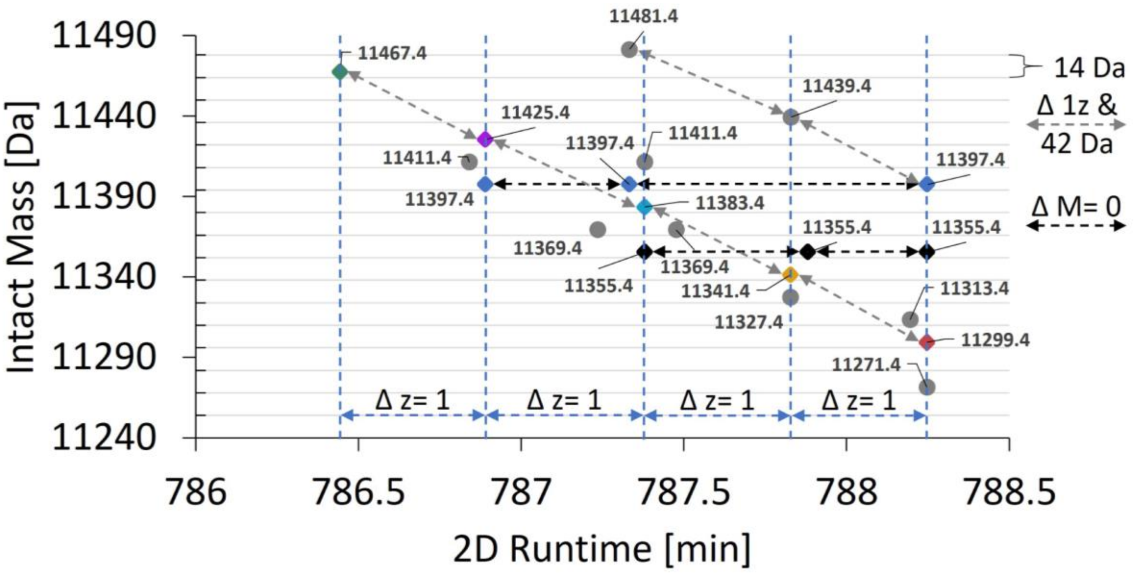

*Figure S 5: Intact mass vs. 2D runtime of selected proteoforms attributed to histone H4, based on the MS1 level. Diamonds represent proteoforms shown in Figure 5 D, black arrows indicate isobaric proteoforms, and grey arrows indicate proteoforms varying in charge from Figure 5 B. Blue arrows and dashed lines indicate distinct groups that are separated based on varying charge.*

#### Comparison of Conventional Sheathless nanoLC-MS Coupling Versus Sheath Liquid-Based nanoLC-MS Coupling

In order to evaluate the influence of the sheath liquid interface on dilution and ionization, additional nanoLC measurements were carried out. For the comparison, a sample was measured using identical nanoLC conditions but two different interfaces. The measurement using the standard nanoLC-MS interface was performed as described in the experimental section using a 20 µm ID 10 µm tip emitter from CoAnn Technologies without any sheath liquid. For the nanoLC-nanoCEasy-MS measurements, the 20µm ID 10 µm tip emitter from CoAnn Technologies was replaced by a 20 µm ID capillary with the same length as the emitter obtained from polymicro which was etched using hydrofluoric acid and placed in the nanoCEasy interface in the same way as the CZE capillary was etched and placed in the nanoCEasy interface (more information about etching, coupling of CZE capillaries with the nanoCEasy interface, and operation of the interface can be found in the materials and methods chapter). Except for the ESI voltage, identical MS methods were used. ESI voltage for standard nanoLC-MS was 1800 V, while the nanoCEasy interface was operated at 2000 V. Figure S 6 shows the difference in intensity for the two used interfaces. The >10-fold intensity difference is assumed to be based primarily on dilution of the nanoLC effluent with sheath liquid.

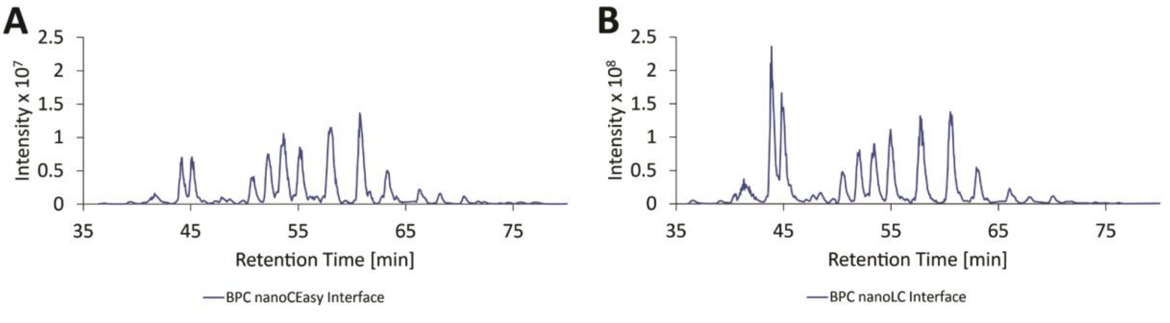

*Figure S 6: nanoLC measurements of Caucasian colon adenocarcinoma cells (CaCo-2) after sample preparation. A) BPC after deconvolution obtained while coupling the nanoLC to the Orbitrap mass spectrometer using the nanoCEasy interface. B) BPC after deconvolution obtained while coupling the nanoLC to the Orbitrap mass spectrometer using the standard interface described in the experimental section.*
